# Identification and characterisation of serotonin signalling in the potato cyst nematode *Globodera pallida* reveals new targets for crop protection

**DOI:** 10.1101/358358

**Authors:** Anna Crisford, Fernando Calahorro, Elizabeth Ludlow, Jessica M.C. Marvin, Jennifer K. Hibbard, Catherine J. Lilley, James Kearn, Francesca Keefe, Rachael Harmer, Peter E. Urwin, Vincent O’Connor, Lindy Holden-Dye

## Abstract

Plant parasitic nematodes are microscopic pests that invade plant roots and cause extensive damage to crops worldwide. To investigate mechanisms underpinning their parasitic behaviour we used a chemical biology approach: We discovered that reserpine, a plant alkaloid known for its antagonism of the mammalian vesicular monoamine transporter VMAT and ability to impart a global depletion of synaptic biogenic amines in the nervous system, potently impairs the ability of the potato cyst nematode *Globodera pallida* to enter the host plant root. We show that this effect of reserpine is mediated by an inhibition of serotonergic signalling that is essential for activation of the stylet, a lance-like organ that protrudes from the mouth of the worm and which is used to pierce the host root to gain access. Prompted by this we identified core molecular components of *G. pallida* serotonin signalling encompassing the target of reserpine, VMAT; the synthetic enzyme for serotonin, tryptophan hydroxylase; the G protein coupled receptor SER-7 and the serotonin-gated chloride channel MOD-1. We found that inhibitors of tryptophan hydroxylase, SER-7 and MOD-1 phenocopy the plant protecting action of reserpine. Thus targeting the serotonin signalling pathway presents a promising new route to control plant parasitic nematodes.

**Summary:** Indian snakeroot, an herbal medicine prepared from the roots of the shrub *Rauwolfia serpentina*, has been used for centuries for its calming action. The major active constituent is reserpine which works by depleting a specific class of mood regulating chemical in the brain, the biogenic amines. We have discovered a remarkable effect of reserpine on a pest of global concern, the plant parasitic nematodes. These microscopic worms invade the roots of crops presenting a severe threat to food production. We show that reserpine disables serotonin signalling in the worm’s ‘brain’ that regulates the rhythmic thrusting of the stylet: a lance-like structure that protrudes from its mouth to pierce the plant root and which is essential to its parasitic lifecycle. Thus, reserpine joins nicotine as another intriguing example of Nature evolving its own protection against pests. We have identified key components of the serotonin signalling pathway in the potato cyst nematode *Globodera pallida* and show that chemicals that target these sites inhibit the ability of the nematode to invade its host plant. We conclude that biogenic amine transmitters are intimately involved in the worm’s parasitic behaviour and provide a new discrete route to crop protection.

## Introduction

Plant parasitic nematodes (PPNs) are microscopic worms that invade the roots of plants causing $125 billion of crop damage per annum [1]. Chemicals deployed to protect crops from PPNs are typically either nematostatics which paralyse the worms by acting as cholinesterase inhibitors or are metabolic poisons [2]. However, the off-target toxicity of these agents is proving unacceptable and they are being removed from use (e.g. EU regulation EC 1107/2009). This presents a growing economic burden that demands new approaches to crop protection. One of the ways to interfere with the infectivity of PPNs is to selectively and discretely disable behaviours that are intrinsic to their parasitic life cycle. Here we provide evidence that serotonin signalling plays an important role in the parasitic life cycle of a PPN and that this provides new targets to protect the host plant.

The subject of this investigation is the sedentary endoparasitic potato cyst nematode, of which there are two major, closely related species *Globodera pallida* and *Globodera rostochiensis*. Together these are important crop pests of worldwide economic significance [3,4]. The particular focus of this work is the white potato cyst nematode *G. pallida*, for which there is no single, dominant potato natural resistance gene available. It has a complex life cycle [5]: The parasitic cycle starts when second-stage juveniles (J2s) hatch from eggs and emerge from the cysts in close proximity to the host roots. These non-feeding juveniles are constrained by their energy stores and have limited time to locate and infect host plant roots. Once the roots are located J2s penetrate an epidermal cell, generally in the zone of elongation, and migrate intracellularly inside the roots to establish a feeding site. Feeding J2s progress through subsequent developmental stages with the vermiform adult males leaving the roots and adult females forming a cyst with hundreds of eggs enclosed within their tanned body wall. The plant completes its cycle and the cysts are released into the soil where new worms will emerge in response to signals from suitable host roots in the next potato cropping season. This cycle can be repeated with a delay of 20 to 30 years, as the eggs are protected and remain dormant in the cysts until favourable conditions for host plant invasion are established. Thus, once a field is contaminated with *Globodera spp.* the threat to crops can be very enduring.

Disruption of the infectivity cycle at the earliest points possible is predicted to lead not only to a reduction in established nematodes and subsequent population build-up but, importantly, could prevent the root damage associated with the early destructive migration of J2s. This damage can reduce the rate of root growth and decrease rates of uptake for water and nutrients [6] likely contributing to reduced tolerance even amongst some resistant potato cultivars. The early target phase encompasses hatching, directed locomotion towards the host plant, detection of the host plant root and root invasion. An important component of these behaviours is the ability of the worm to appropriately activate its stylet, a hollow lance-like structure that can protrude from the mouth of the worm, and is implicated in both hatching and penetration of the host plant root [7,8]. Once inside the host root, secretions from the nematode pharyngeal glands are introduced via the stylet into a selected plant cell. These secretions are intimately involved in establishing and maintaining the syncytial feeding site from which the stylet itself then provides the sole route for nutrient acquisition. All of these behaviours are the output of the nematode’s nervous system but little is currently understood of their neurobiology [8].

Here we report a remarkable effect of reserpine, which impairs the ability of *G. pallida* J2s to invade the host plant root. Reserpine is a naturally occurring plant alkaloid from the shrub *Rauwolfia serpentina*. We show reserpine protects roots from *G. pallida* invasion through a potent inhibition of the vesicular monoamine transporter, VMAT, that impacts on the activity of the J2’s stylet required for root invasion. This prompted us to identify further core molecular components of the serotonin signalling pathway in *G. pallida* and we report that targeting these with chemical inhibitors can also protect roots from infection. Given the universally important role of the stylet to the life cycle and feeding of PPNs we argue that targeting serotonin signalling pathways presents an under-utilised and promising new route to control PPNs.

## Results

### The inhibitory effect of reserpine on G. pallida root invasion behaviours

Pre-incubation of *G. pallida* J2s with 100 µM reserpine prior to their inoculation onto potato hairy roots in tissue culture significantly decreased the number of nematodes present in the roots 13 days later (Fig 1A). In order to invade the host root the J2s must be motile and capable of thrusting their stylet in a rhythmic manner to pierce the plant cell walls [5]. We tested the effect of reserpine on *G. pallida* motility in a dispersal assay in which J2s were placed in the centre of an agar arena and the number of worms present at the origin after a given period of time was scored. Dispersal was significantly reduced after pre-incubation with reserpine (50 µM) for 18 h (Fig 1B) consistent with previous reports that monoaminergic transmission, specifically that involving serotonin, is required for motility of plant parasitic nematodes [8,9]. Taken together, the data are consistent with the interpretation that reserpine depletes endogenous serotonin in the circuit that is required to stimulate dispersal (Fig 2A), an effect that would contribute to the impairment of host plant invasion.

**Figure 1.**
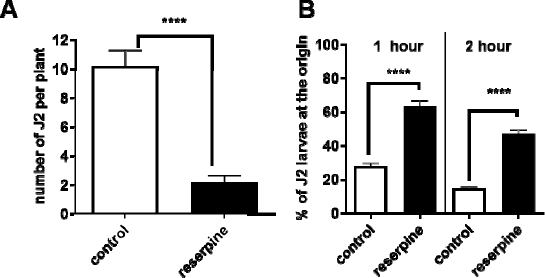
Reserpine inhibits *G. pallida* host plant invasion behaviour. **A.** J2s were collected at 24 h post hatching and pre-incubated with 24 h in water without (control), or with the addition of reserpine (100 µM). J2s were applied to individual potato hairy root cultures at 3 infection points with 25 J2s per infection point. 13 days later roots were stained with acid fuchsin and parasitic nematodes visualised. Data are mean ± s.e.m. for 20 plants; **** p<0.0001 unpaired Student’s t-test. **B.** The effect of reserpine on J2 motili was tested in a dispersal assay in which their ability to move away from the central point of an agar arena was determined. For each assay about 50 to 100 J2s were presoaked in water (control), or water with 50 µM reserpine for 18 h. They were then pipetted onto the centre of the plate in a minimum (˜ 5 µL) volume of liquid. The percentage of J2s remaining at the origin was determined after 1 and 2 h. Data are mean ± s.e.m. of 4 plates for each experimental group. One way ANOVA with Bonferroni’s multiple comparisons; ****p<0.001.

**Figure 2.**
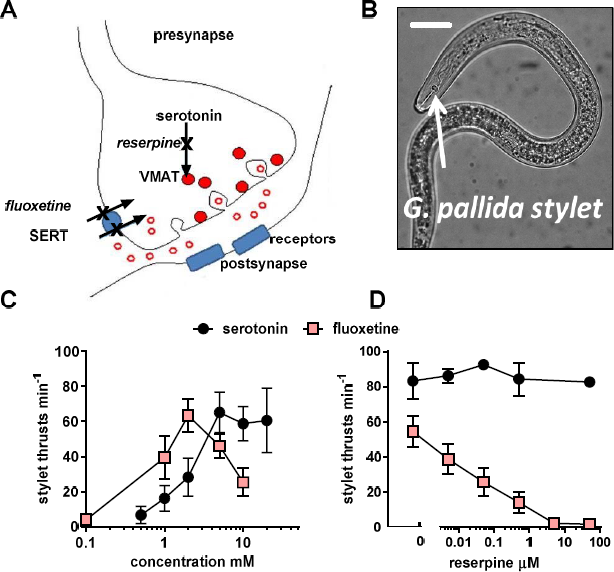
**Reserpine inhibits stylet thrusting triggered by endogenous, but not exogenous, serotonin. A.** The cartoon depicts a serotonergic synapse, with the presynapse as the site of serotonin synthesis and release and the postsynapse which harbours serotonin receptors. In mammalian systems reserpine acts presynaptically to deplete the storage of the neurotransmitter serotonin by preventing its uptake into vesicles (shown as red circles) by an inhibition of the vesicular monoamine transporter, VMAT [34]. This results in a lack of serotonin presynaptically and a block of serotonergic neurotransmission. Fluoxetine (Prozac^®^) blocks the plasma membrane serotonin transporter, SERT. Thus it increases synaptic levels of serotonin (shown by open red circles) by preventing its reuptake into the presynaptic terminal following its release. In this way fluoxetine can act indirectly to stimulate transmission at the serotonergic synapse. This effect of fluoxetine is susceptible to block by reserpine as it requires endogenous serotonin to elicit its effect. However, the response to exogenous serotonin circumvents reserpine inhibition as it acts directly on the postsynaptic receptors. **B**. The *G. pallida* stylet is a lance-like structure that can be thrust out of the mouth of the worm (in this image it is shown in the retracted position) in a rhythmic manner to initiate hatching and root invasion. Inside the root it is used for migration and to support feeding and interaction with the host. The activity of the stylet can be visually scored by counting the number of thrusts made in 1 min. Scale bar ˜ 20 µm. **C.** J2s were incubated in either serotonin or fluoxetine at the concentrations indicated for 1 h and then the number of stylet thrusts made in 1 min was counted. Data are mean ± s.e.m.; n= 8 to 17 for each time point. **D.** Reserpine blocked the stylet response to fluoxetine but not serotonin. J2s were presoaked in reserpine at the concentration indicated for 24 h and subsequently immersed in either 10 mM serotonin or 2 mM fluoxetine for 30 min and stylet thrusting scored for 1 min. Data are mean ± s.e.m.; n=10 J2s for each data point.

Furthermore, serotonin is a known activator of stylet thrusting [9]. We therefore speculated that reserpine may also impair this behaviour which contributes to root invasion. Fluoxetine, more commonly known as the antidepressant Prozac^®^, was deployed to activate stylet thrusting of J2s *in vitro* so that the inhibitory action of reserpine could be tested [10]. This compound blocks the synaptic plasma membrane serotonin transporter, and by preventing re-uptake of serotonin, increases its concentration in the synaptic cleft which in turn activates the postsynaptic receptors (Fig 2A). Stylet thrusting can be visually scored by counting the frequency of projection/retraction cycles of the stylet (Fig 2B): Both serotonin and fluoxetine elicited a time-dependent and concentration-dependent stimulation of stylet thrusting. Responses reached a plateau after 30 min exposure (stylet thrusts for 2 mM fluoxetine after 30 min = 50 ± 11 min^−1^, compared to 58 ± 9 min^−1^ after 1 h, p>0.05: Stylet thrusts for 10 mM serotonin after 30 min = 54 ± 10 min^−1^, compared to 58 ± 10 min^−1^ after 1h, n=17, p >0.05). The concentration–response curve for serotonin after 1 h reached a maximum plateau above 2 mM whilst fluoxetine had a bell-shaped concentration response curve (Fig 2C). This may indicate additional pharmacological sites of action for fluoxetine over and above the plasma membrane transporter for serotonin, most likely serotonin receptors. Serotonin receptor antagonism by fluoxetine at higher concentrations has been reported previously for *C. elegans* [11]. We tested the effect of reserpine against the maximally effective concentration of serotonin (10 mM) and fluoxetine (2 mM): Reserpine potently blocked the stylet response to fluoxetine but not the response to serotonin (Fig 2D) consistent with an interpretation in which the response to fluoxetine requires the presence of correctly stored vesicular serotonin which is depleted by the VMAT-blocking action of reserpine (see Fig 2A). We tested the reversibility of this reserpine inhibition by exposing *G. pallida* J2s to 50 µM reserpine with 2 mM fluoxetine for 10 h and then transferring them to reserpine free solution, with 2 mM fluoxetine, the ‘recovery’ period. Controls consisted of J2s that were exposed to 2 mM fluoxetine only. After 22 h in the recovery period stylet thrusting in the control group was 62.3 ± 6.5 min-1(n=10) whilst in the reserpine treated group it was 0.2 ± 0.2 (n=5; p<0.001). Thus the inhibition of stylet thrusting by reserpine is sustained following its removal.

Together these data suggest that reserpine treatment may lead to a sustained depletion of endogenous serotonin in neural circuits regulating motility and stylet thrusting leading to an inability of the nematode to invade the host root. Furthermore, it indicates that serotonin signalling plays a key role in host plant invasion behaviour and therefore we embarked on a characterisation of core components of serotonergic neurotransmission with a view to testing these as potential targets for crop protection.

### *G. pallida* VMAT is a functional orthologue of *C. elegans* CAT-1

We identified the molecular target for reserpine in *G. pallida* using the sequence of the *C. elegans* vesicular monoamine transporter *cat-1* (WBGene00000295) which is well characterised [12] and for which there are two isoforms, *cat-1a* and *cat-1b*. We mined the *G. pallida* draft genome assembly (4) and identified a single *cat-1* homologue (GPLIN_000654600) which we designated *gpa-cat-1*. We cloned two variants of *G. pallida cat-1* from *J2* cDNA, which had 2 and 3 amino acid differences, respectively, from the published sequence. These amino acid changes could not be assigned to any known functions of CAT-1 domains in *C. elegans* and are thus likely to be allelic variants. (Supplementary Fig 1).

In adult hermaphrodite *C. elegans cat-1* is expressed in all serotonergic and dopaminergic neurons [13]. *C. elegans cat-1(ok411)* functional null mutants exhibit a distinctive feeding phenotype; they cannot sustain fast pharyngeal pumping on food [12]. This reflects the major role that serotonin has in stimulating pharyngeal pumping in the presence of food [14-16]. To test whether or not the sequence identified in *G. pallida* is a functional orthologue of *ce-cat-1* we expressed the *gpa-cat-1* cDNA in, the null mutant *C. elegans* and tested for rescue of the *C. elegans* pharyngeal phenotype. The two cloned allelic variants of *gpa-cat-1* were both individually expressed in *C. elegans cat-1(ok411)* from a pan-neuronal promoter *psnb-1* [17] and transgenic lines scored for the *cat-1 (ok411)* pharyngeal phenotype. This was compared to rescue achieved by expression of *C. elegans cat-1* from the same promoter. As *C. elegans cat-1a* is more similar to *G. pallida cat-1* this was used for the control rescue experiments. Transformed worms were identified by co-expression of *gfp* from the *myo-3* promoter which provides readily identifiable fluorescence in the body wall muscle of transgenic lines. The control worms expressing *myo-3::gfp* had the same pumping rate as untransformed *cat-1* mutants (*myo-3::gfp* 182 ± 2 and *cat-1* 192 ± 2 pumps min^−1^, respectively; p>0.05) indicating expression of the transformation marker alone does not impact on pharyngeal pumping. Both *gpa-cat-1* and *C. elegans cat-1a* rescued the pharyngeal pumping phenotype of *cat-1(ok411)* mutant worms to the levels of wild type (Fig 3A). Thus *gpa-cat-1* encodes a vesicular monoamine transporter functional in *C. elegans*. The same rescue was observed using either *gpa-cat-1a* or *gpa-cat-1b* supporting the conclusion that both these sequences encode a functional VMAT in *G. pallida*.

**Figure 3.**
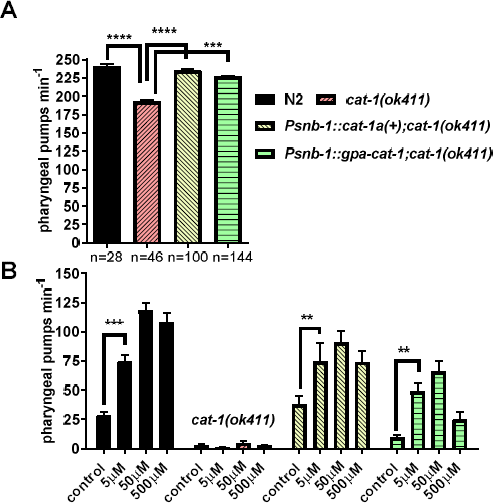
**Functional characterisation of a gene encoding** ***G. pallida*** **VMAT,** ***gpa-cat-1*****. A.** Expression of *G. pallida cat-1* (*gpa-cat-1)* rescues the pharyngeal phenotype of *C. elegans cat-1(ok411)*. A comparison of the pumping rate on food for N2, *cat-1(ok411)* and transgenic lines of *cat-1(ok411)* expressing either wild type *C. elegans cat-1*, *cat-1(+)* or *gpa-cat-1* behind a pan-neuronal promoter (*Psnb-1*). Four stable lines for each transgenic strain were tested and the data are pooled for presentation. Data are mean ± s.e.m; ‘n’ is shown in brackets. *** p<0.001; **** p<0.0001; One way ANOVA with Bonferroni’s multiple comparisons. **B**. Expression of *gpa-cat-1* in *C. elegans cat-1(ok411)* reinstated sensitivity to fluoxetine. One day old hermaphrodite *C. elegans* were placed on agar plates that either had no drug, ’control’, or had been prepared with fluoxetine (5 to 500 µM). After 1 h the rate of pharyngeal pumping was scored in each worm for 1 min. N2 wild type worms pumped at a low rate which increased in a concentration-dependent manner in the presence of fluoxetine. Fluoxetine did not stimulate pumping in *cat-1(ok411)* but the response was restored in the transgenic worms expressing either *cat-1(+)* or *gpa-cat-1* behind a pan-neuronal promoter (*psnb-1*). Two stable lines for each transgenic strain were tested and the data are pooled. Data are the mean ± s.e.m.; n ≥ 17; **p<0.01; *** p<0.001; Two way ANOVA with Bonferroni’s multiple comparisons. For the sake of clarity only one comparison is shown for each strain between control and 5 µM fluoxetine.

The function of CAT-1 may also be assessed through the pharmacological response of the worms to fluoxetine. The impact of fluoxetine on pharyngeal pumping is not apparent in well fed *C. elegans* in the presence of food (bacteria) as the worms’ pharyngeal activity is maximally activated by the presence of the bacteria. However, in the absence of bacteria pharyngeal activity is much reduced: Under these conditions pharmacological stimulation of pharyngeal pumping with fluoxetine, which elevates synaptic serotonin, a key signal for activation of pumping [16,18], can be observed. The stimulatory effect of fluoxetine on *C. elegans* pharyngeal pumping in the absence of food was maximal after 1 h exposure (data not shown) and concentration-dependent (Fig 3B). Similar stimulation was not observed in the *C. elegans cat-1* mutant, consistent with a model in which fluoxetine acts by elevating endogenous synaptic levels of serotonin by blocking reuptake via the serotonin transporter, SERT (Fig 2A)[19]. Higher concentrations of fluoxetine (1 mM and 2 mM) or longer exposure times at 500 µM, did not cause a stimulation of pharyngeal pumping which probably reflects that fact that at higher concentrations fluoxetine is an antagonist of serotonin receptors [11,15] thus any stimulatory action would be inhibited by concomitant receptor blockade. Importantly, with respect to establishing the function of *gpa-cat-1*, the low dose stimulatory effect of fluoxetine on pharyngeal pumping that was absent in the *C. elegans cat-1* mutant was restored by expression of either wild-type *C. elegans cat-1* or *gpa-cat-1* (Fig 3B). This confirms the identification of the vesicular monoamine transporter, VMAT, from *G. pallida.*

### Cloning and functional analysis of G. pallida tph-1

To further test the role of serotonin in the parasitism of PPNs we investigated the role of the synthetic enzyme for serotonin, tryptophan hydroxylase. The *C. elegans* gene encoding tryptophan hydroxylase *tph-1* (WBGene00006600) has two predicted isoforms, *tph-1a* and *tph-1b*. *C. elegans tph-1a* is assembled from 11 exons, while *tph-1b* has a shorter 5’ terminus and is assembled from 9 exons. The *G. pallida* genome assembly contains a putative orthologue (GPLIN_000790300) that we designated *gpa-tph-1*. A single isoform was cloned from cDNA that more closely resembles the longer *ce-tph-1a* (Supplementary Fig 2).

A *C. elegans* functional null mutant for *tph-1(mg280)* has reduced pharyngeal pumping due to the lack of serotonin [13,20]. To confirm the functional identity of *gpa-tph-1* as a genuine tryptophan hydroxylase we tested the ability of this gene to rescue the *C. elegans tph-1(mg280)* pharyngeal phenotype and compared this to the rescue obtained by expression of *C. elegans tph-1a(+)*. Injection of *tph-1* with the transformation marker, *myo-3::gfp*, alone did not change pumping rate (*myo-3::gfp* 179 ± 4 pumps per minute; *tph-1(mg280)* 169 ± 3 pumps min^−1^; p>0.05). *G. pallida tph-1* rescued the pumping phenotype of the *C. elegans tph-1(mg280)* mutant to the levels of wild type in a similar manner to expression of *C. elegans tph-1* (Fig 4A), supporting the conclusion that the *G. pallida* and *C. elegans tph-1* genes are functional orthologues.

**Figure 4.**
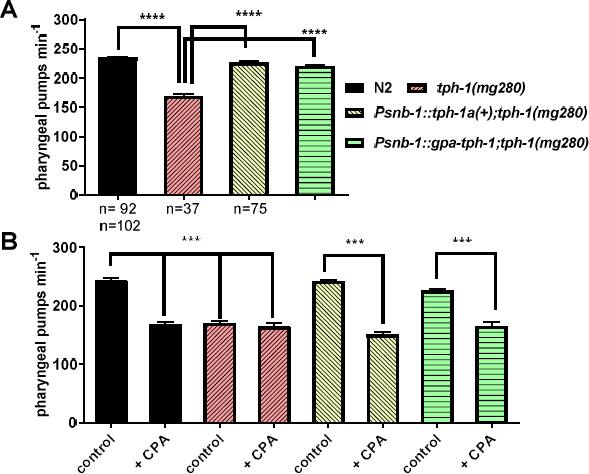
**Functional characterisation of a gene encoding** ***G. pallida*** **tryptophan hydroxylase,** ***gpa-tph-1.*** **A.** Expression of *gpa-tph-1* rescued the pharyngeal phenotype of *C. elegans tph-1(mg280)* similar to expression of a wild type copy of *C. elegans tph-1*, *tph-1(+)*. Both genes were expressed from the pan-neuronal promoter *Psnb-1*. Pharyngeal pumping rate of one day old adult hermaphrodites on food was scored for 1 min for each worm. Four stable lines for each transgenic strain were tested and the data are pooled. Data are mean ± s.e.m; ‘n’ is indicated on the graph; **** p<0.0001; one way ANOVA with Bonferroni’s multiple comparisons. **B.** Expression of either *C. elegans tph-1(+)* or *gpa-tph-1* in the *C. elegans* mutant *tph-1(mg280)* conferred sensitivity to the tryptophan hydroxylase inhibitor, CPA. One day old adult hermaphrodites were incubated for 2 h on agar plates with food and without, (control), or with 1 mM CPA. Two stable lines were tested for each transgenic strain and the data are pooled. Data are the mean ± SEM of n ≥ 10; ***p<0.001; one way ANOVA with Bonferroni’s multiple comparisons.

We next wanted to interrogate the functional importance of *G. pallida* TPH-1 in root invasion. Unfortunately, RNA interference through soaking in double-stranded (ds)RNA has non-specific toxic effects on *G. pallida* confounding the use of this experimental approach [21]. Therefore we investigated pharmacological blockers as a route to establish the functional involvement of serotonin signalling in *G. pallida* parasitism. We tested whether 4-chloro-DL-phenylalanine methyl ester hydrochloride (CPA), which is an inhibitor of the mammalian tryptophan hydroxylase [22], could be used as a chemical tool to test the function of TPH in*G. pallida.* To achieve this we first established the ability of CPA to block both *C. elegans* and *G. pallida* TPH-1 expressed in the *C. elegans tph-1* mutant. Indeed, we found that pharyngeal pumping of wild type *C. elegans* was significantly inhibited by CPA (10 mM) to a level that phenocopied the *tph-1* mutant and furthermore exposure of the mutant to CPA did not cause any further reduction in pharyngeal pumping (Fig 4B). The ability of CPA to inhibit pharyngeal pumping was restored in *tph-1* mutants by expression of either *C. elegans* or *G. pallida tph-1* (Fig 6B). These data are consistent with CPA acting as a selective inhibitor of *C. elegans* and *G. pallida* TPH-1.

### Targeting TPH-1 impairs G. pallida root invasion behaviour

We used CPA to test the role of the enzyme encoded by *gpa-tph-1* in root invasion behaviour of *G. pallida*. *G. pallida* J2s were treated with 10 mM CPA overnight after which treated and control worms were soaked in serotonin (10 mM) or fluoxetine (2 mM) for 30 min and stylet thrusting visually scored. The worms incubated in CPA overnight had reduced stylet thrusting in response to fluoxetine but maintained their response to serotonin suggesting that CPA inhibited TPH-1 enzyme activity and thus serotonin synthesis (Fig 5A) and consequently the response to fluoxetine which is dependent on endogenous serotonin. Treatment of J2s with CPA (100 µM) prior to infection significantly reduced the number of nematodes present in potato roots 13 days after inoculation (Fig 5B) adding further evidence to support a role for serotonin signalling in root invasion.

**Figure 5.**
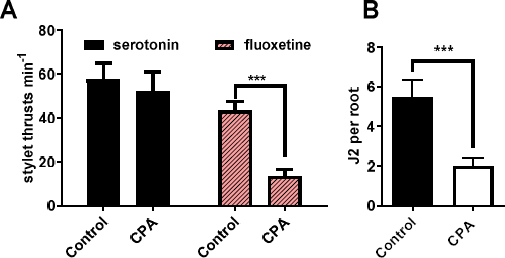
**The role of TPH-1 in host plant invasion behaviour. A.** *G. pallida* J2s were soaked in either water (control) or water with the TPH inhibitor CPA (100 µM) for 24 h. This was followed by the addition of either serotonin (10 mM) or fluoxetine (2 mM) for 30 min and stylet thrusting was scored for 1 min. Data are mean ± s.e.m. n = 27 for serotonin and n=40 for fluoxetine. *** p<0.001; Two way ANOVA with Bonferroni’s multiple comparisons. **B.** CPA impaired the ability of J2s in root invasion. J2s were collected at 24 h post hatching and pre-incubated for 24 h in water without (control), or with the addition of CPA (100 µM). J2s were applied to individual potato hairy root cultures at 3 infection points with 25 J2s per infection point. 13 days later roots were stained with acid fuchsin and nematodes visualised. Data are mean ± s.e.m. for 19 plants; **** p<0.001 unpaired Student’s t-test.

### Functional and pharmacological characterisation of *G. pallida* serotonin receptors

The characterisation of *G. pallida* VMAT and TPH described above strongly supports a role for serotonergic transmission in host root invasion and provides a justification for pursuing the identify of *G. pallida* serotonin receptors that are involved in this behaviour. For this we used the well characterized serotonergic signalling pathway in *C. elegans* as our reference point [23-27]. Similar to their closest homologues in humans, *C. elegans* SER-1 and SER-4 have a low affinity for serotonin whilst SER-7 has a high affinity. Interestingly, MOD-1, which has a high affinity for serotonin, is a serotonin-gated chloride channel which is selectively found in the invertebrate phyla [28]. *G. pallida* putative orthologues of the *C. elegans* genes encoding serotonin receptors were identified by reciprocal best BLAST hit analysis of the *G. pallida* predicted gene complement. This identified *G. pallida* genes encoding candidate orthologues for the *C. elegans* receptors SER-4, SER-7 and MOD-1 although clear orthologues of SER-1 and SER-5 were not found (Table 1). In this study we focused on characterisation of SER-7, selected because of its key role in regulating feeding behaviour in *C. elegans* [15] and MOD-1, selected because of its important role in regulating locomotion underpinning exploration [29] in the anticipation that these receptors may subserve similar behaviours in plant parasitic nematodes.

**Table 1.** Identification of *C. elegans* serotonin receptor homologues in *G. pallida*. *G. pallida* homologues of the five *C. elegans* genes encoding serotonin receptors were identified by reciprocal best BLAST hit analysis of the *G. pallida* predicted gene complement[3].

| *C. elegans* receptor | Isoforms in *C. elegans* | Potential orthologue in *G. pallida* | Closest *G. pallida* match<br>Amino acid identity (%) to *C. elegans* | Protein function |
| --- | --- | --- | --- | --- |
| SER-1 | *ser-1a*<br>*ser-1b* | NO | 18.8 | G-protein coupled receptor |
| SER-4 |  | YES | 41.2 | G-protein coupled 5-HT receptor |
| SER-5 |  | ? |  | G-protein coupled receptor |
| SER-7 | *ser-7a*<br>*ser-7b*<br>*ser-7c* | YES | 35 | G-protein coupled receptor |
| MOD-1 | *mod-1a*<br>*mod-1b*<br>*mod-1c* | YES | 56.8 | Ligand- gated chloride channel |

SER-7 is a key determinant of *C. elegans* pharyngeal pumping [15,18,25]. This is demonstrated by the observation that *C. elegans* null mutants for *ser-7* display irregular pumping on food and have a reduced response to serotonin-stimulated pumping in the absence of food [15]. We cloned the closest homologue of *C. elegans ser-7* from *G. pallida* (Supplementary Fig 3). To determine if this *G. pallida* sequence encodes a *bona fide* serotonin receptor, we tested its ability to rescue the lack of serotonin- stimulated pharyngeal pumping in the *C. elegans ser-7* mutant. We found that expression of either the putative *G. pallida ser-7* or *C. elegans ser-7* from the pan-neuronal promoter *snb-1* restored sensitivity of the pharyngeal system to serotonin (Fig 6A). In these *in vivo* whole organism experiments a high concentration of serotonin is required to activate the pharynx as the worm’s cuticle presents a permeability barrier to drug access [30] confounding discrete pharmacological analysis of the SER-7 receptor. Moreover, although SER-7 is an important contributor to the pharyngeal serotonin response in the intact worm it is not the sole receptor involved, as evidenced by the residual response to serotonin in the *ser-7* mutant (Fig 6A). To circumvent these confounds we analysed the pharyngeal response to serotonin in a cut-head assay in which the intact pharynx is carefully cut from the rest of the worm (Fig 6B). This enables drugs to access the tissue without having to cross the nematode cuticle and permits more precise interrogation of the concentration-dependence of the compounds’ effects on pharyngeal activity [16,31]. Furthermore, the response of the pharynx in the cut-head preparation to serotonin is entirely dependent on SER-7 (Fig 6C). The isolated pharynx has previously been shown to be 2 to 3 orders of magnitude more sensitive to the stimulatory action of serotonin compared to the intact worm [16]. Consistent with this, wild type worms showed a concentration-dependent increase in pharyngeal pumping in response to serotonin (EC_50_ 223 nM, 95% confidence from 164 nM to 304 nM) whilst s*er-7(tm1325)* mutants did not respond even to 100 µM serotonin (Fig 6C). This is in agreement with earlier studies showing SER-7 is the key receptor intrinsic to the pharyngeal system that mediates serotonin activation of feeding [15]. Pharyngeal pumping in transgenic worms expressing either *psnb-1::ser-7a(+)* or *psnb-1::gpa-ser-7* was activated by serotonin with an EC_50_ of 782 nM (95% confidence from 457nM to 1.34 µM) and EC 50 3.9 µM (95% confidence from 2.16 µM to 7.08 µM), respectively (Fig 6C). We used this cut head pharmacological assay to test a putative antagonist of the SER-7 receptor, methiothepin [25] and showed that this compound blocked the stimulatory effect of serotonin in transgenics expressing either *C. elegans ser-7(+)* or *G. pallida ser-7* (Fig 6D) with around 50% inhibition at 100 nM. Together, this supports the conclusion that *gpa-ser-7* encodes a *G. pallida* serotonin receptor which has the closest similarity to *C. elegans* SER-7.

**Figure 6.**
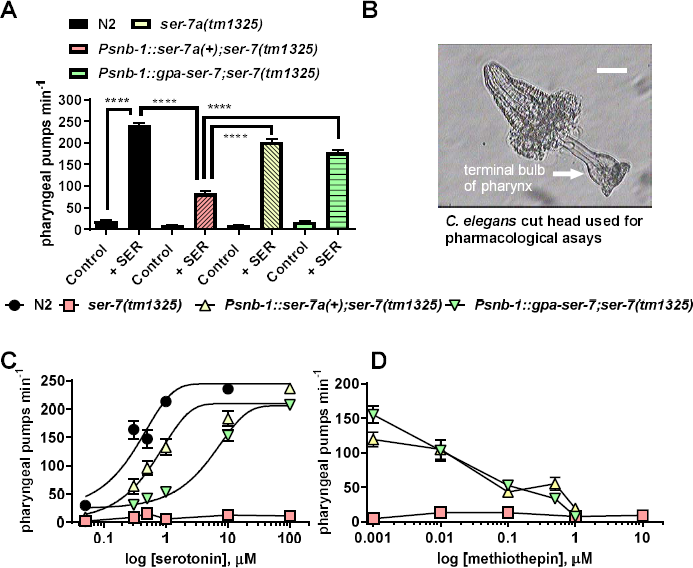
**Characterisation of** ***G. pallida*** **SER-7 A.** Expression of *gpa-ser-7* rescued the pharyngeal phenotype of *C. elegans ser-7(tm1325)* similar to expression of a wild type copy of *C. elegans ser-7a*, *ser-7a(+)*. Both genes were expressed from the pan-neuronal promoter *Psnb-1*. Pharyngeal pumping rate of one day old adult hermaphrodites after 20 min off food was scored for 1 min for each worm either in the absence (control) or presence (SER) of 10 mM serotonin. Three stable lines for each transgenic strain were tested and the data are pooled. Data are mean ± s.e.m; n ≥ 20; **** p<0.0001; one way ANOVA with Bonferroni’s multiple comparisons. Note that the whilst the pumping rate of *ser-7(tm1325)* was increased by 10 mM serotonin this response was significantly lower than that for wild type and both transgenic strains. **B.** An image of the cut head preparation that was used for the experiments shown in C and D. Cutting the head exposes the pharynx to the external solution allowing ready access of applied drugs to the receptors regulating the pharyngeal network. Pharyngeal pumps were recorded visually by counting the contraction-relaxation cycles of the terminal bulb. Scale bar ˜25 µm. **C.** Concentration-response of the pumping rate of the cut head pharyngeal preparations to serotonin for wild type (N2), *ser-7(tm1325)* and transgenic lines of *ser-7(tm1325)* expressing either *C. elegans* or *G. pallida ser-7* behind a pan-neuronal promoter (*psnb-1*). Note that in the dissected pharyngeal preparation *ser-7(tm1325)* mutants do not respond to even 100 µM serotonin. Data are the mean ± s.e.m.; n ≥ 20. **D.** Methiothepin blocked the response to serotonin in both transgenic strains. Cut heads were pre-incubated with methiothepin at the concentrations indicated for 5 min after which 100 µM serotonin was added and after a further 10 min the pumping rate was scored for 1 min. Two stable lines for each transgenic strain were tested and the data are pooled. Data are the mean ± s.e.m; n ≥ 5.

We also cloned a putative orthologue of the *C. elegans* serotonin-gated chloride channel *mod-1* from *G. pallida* (Supplementary Fig 4). This receptor belongs to a family of biogenic amine-gated chloride channels which are unique to the invertebrate phyla and thus of particular interest with respect to the development of novel chemical control agents with selective toxicity towards plant parasitic nematodes.

We found that a null mutant for *C. elegans mod-1, (ok103),* has a pharyngeal phenotype with a slightly reduced rate of pharyngeal pumping compared to controls. The pumping rate on food for *mod-1(ok103)* was 214 ± 3 min^−1^ (n=10) and in the control injected line (*myo-3::gfp;mod-1(ok103)*) 189 ± 7 min^−1^ (n=12) compared to 252 ± 4 (n=7) in wild-type (p<0.0001, one way ANOVA with Bonferroni’s multiple comparisons). This effect on pharyngeal pumping was rescued in lines expressing *C. elegans mod-1(+)* from the pan-neuronal promoter *Psnb-1* (230 ± 7 min^−1^; n=9; p<0.0001). This is consistent with a role for MOD-1 in regulating *C. elegans* feeding behaviour. However, a more distinctive phenotype of *C. elegans mod-1 (ok103)* is that it does not exhibit the acute transient paralysis induced in wild type by a high concentration (33 mM) of serotonin [27]. We found that the putative *G. pallida* sequence encoding the orthologue of the *C. elegans* MOD-1 receptor could completely rescue this serotonin-induced phenotype in *ok103* as robustly as the expression of a wild type *C. elegans mod-1* (Fig 7A). For these experiments we cloned the putative native *C. elegans mod-1* promoter to drive expression of both the *G. pallida* and *C. elegans* sequences. Like SER-7, *C. elegans* MOD-1 is also blocked by methiothepin therefore to further characterise the *G.pallida* receptor we compared the ability of methiothepin to block *C. elegans* and *G. pallida* MOD-1. We showed that the *mod-1* dependent serotonin-induced paralysis in both wild type worms and in worms expressing *G. pallida mod-1* was blocked by methiothepin (Fig 7B). This supports the conclusion that the *G. pallida* sequence encodes a functional orthologue of the *C. elegans* serotonin-gated chloride channel MOD-1.

**Figure 7.**
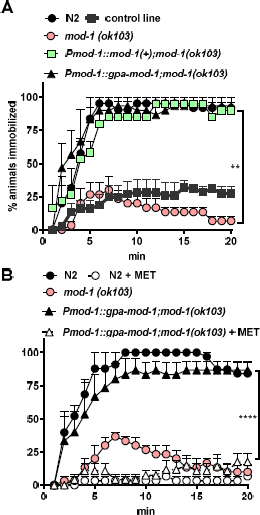
**Characterisation of** ***G. pallida*** **MOD-1. A.** Expression of *gpa-mod-1* rescued the motility phenotype of *C. elegans mod-1(ok103)* similar to expression of a wild type copy of *C. elegans mod-1*, *mod-1(+)*. Both genes were expressed from the putative native *mod-1* promoter*, Pmod-1*. Approximately 10 one day old adult hermaphrodite *C. elegans* were dispensed into each well of a 96-well microtitre plate. At time 0, 33 mM serotonin dissolved in M9 buffer was added to each well and the percentage of immobilised worms in each well was scored every min for up to 20 min. Three stable lines for each transgenic strain were tested and the data are pooled. Data are mean ± s.e.m; n =3 independent experiments; ** p<0.01; two way ANOVA with Bonferroni’s multiple comparisons. **B.** The serotonin-induced paralysis was blocked by methiothepin (MET) in both wild type (N2) and in the transgenic strain expressing *gpa-mod-1* in *mod-1(ok103).* The experiment was performed as in A with the addition of a preincubation in 10 µM MET on an agar plate seeded with OP50 for 2 h and the inclusion of 10 µM MET with serotonin in the microtitre plate wells. Data are mean ± s.e.m; n=3 independent experiments; ** p<0.01; two way ANOVA with Bonferroni’s multiple comparisons.

In view of the evidence that a low micromolar concentration of methiothepin can block both *G. pallida* SER-7 and MOD-1 we speculated that this compound might provide insight into whether either, or both, of these receptors are involved in host plant invasion behaviour. We found that methiothepin is a remarkably potent antagonist of serotonin-induced stylet thrusting in *G. pallida* J2s (Fig 8A) and blocked root invasion (Fig 8B). Interestingly, stylet thrusting is also implicated in egg hatching [5] and we found that methiothepin potently inhibited the emergence of J2s from eggs (Fig 8C) although the J2s inside the eggs appeared viable (Fig 8D). In contrast, exposure of *G. pallida* cysts to 50 µM reserpine did not inhibit hatching (n=20 cysts).

**Figure 8.**
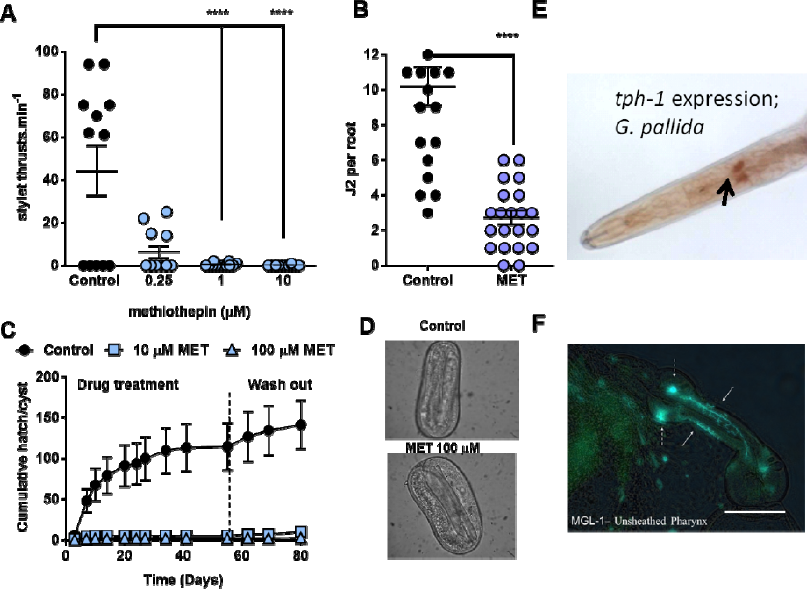
**Methiothepin is a potent inhibitor of*****G. pallida*** **behaviours, hatching and root invasion, that are dependent on serotonin regulated stylet thrusting. A.** *G. pallida* J2s were soaked in either a control solution of 0.5% ethanol or methiothepin at different concentrations for 24 h. They were transferred to 2 mM fluoxetine and after 30 min had elapsed the rate of stylet thrusting was counted for 1 min. Data are mean ± s.e.m.; n=12; p<0.0001; one-way ANOVA with Bonferroni’s multiple comparisons. **B**. Methiothepin (MET) impaired the ability of J2s to invade the host root. J2s were collected at 24 h post hatch and pre-incubated for 24 h in water without (control), or with the addition of MET (100 µM). J2s were applied to individual potato hairy root cultures at 3 infection points with 25 J2s per infection point. 13 days later roots were fuschin stained and visualised for J2. Data are mean ± s.e.m. for 20 plants for control and 21 plants for MET; **** p<0.001 unpaired Student’s t-test. **C.** MET blocked hatching of J2s from cysts. *G. pallida* cysts were soaked in a 24-well plate with 1 cyst per well in the presence PRD modified with either vehicle (PRD), 10 µM methiothepin (M) or 100 µM methiothepin and hatching of J2 juveniles was scored. At 24 days, the cysts were placed into PRD alone to assess recovery of hatching. Methiothepin inhibited hatching and hatching did not recover on removal from drug (P<0.0001). **D.** Representative DIC images of unhatched eggs from cysts treated with a control solution and 100 µM methiothepin. Note that methiothepin treatment does not appear to have negatively impacted the integrity of the unhatched J2 (egg dimensions 120 µm by 60 µm). **E & F.** A comparison of the localization of serotonergic neurons in *G. pallida* (**E**) and *C. elegans* (**F**) suggests that serotonin may subserve similar behaviour functions in these two nematodes. **E.** *In situ* hybridization for *tph-1* reveals two candidate serotonergic neurons in the anterior region of *G. pallida*. **F.** A dissected *C. elegans* pharyngeal preparation showing the location of the two serotonergic neurosecretory motorneurons NSM in the pharyngeal system (*gfp* driven from the promoter *mgl-1* [52]). There are two further extrapharyngeal neurons, ADF, in the anterior (not shown). Together NSM and ADF couple environmental signals reporting the presence of food to behavioural responses of feeding and dwelling [29,53].

In support of a role for serotonin in stylet-mediated behaviours we identified two putative neurons in the anterior nervous system using an *in situ* hybridisation probe targeted at *tph-1* (Fig 8E). These neurons are in a very similar anterior region to the serotonergic neurosecretory motorneurons (NSM) (Fig 8F) and the amphidial neurons ADF in *C. elegans*. Taken together these data are consistent with the interpretation that methiothepin acts as an antagonist of either, or both, SER-7 and MOD-1 to block serotonin signalling in the anterior of the worm that is essential for engaging stylet thrusting in response to environmental stimuli to initiate hatching and root invasion.

## Discussion

The most economically important PPNs are the cyst (e.g. *Heterodera* and *Globodera* spp.) and root-knot nematodes (*Meloidogyne* spp.). These pests present an increasing threat to global food security which is exacerbated by the imposition of restraints on the use of nematicides due to their unacceptable levels of hazard and ecotoxicity. Strategies to improve this situation are highly dependent on achieving a better understanding of the biology underpinning the interaction of these nematodes with their host plants [32,33].

There are discrete differences in the life cycle and behaviour of different species of PPNs but for all of them the ability to regulate their motility and the thrusting behaviour of their stylet in response to discrete environmental cues is core to their parasitic potential. In this study we have improved understanding of the neurobiological basis of these behaviours and shown a critical role for core components of a serotonergic signalling pathway. Our route into this was *via* a chemical biology approach in which we investigated the ability of diagnostic neuroactive drugs to impact on root invasion behaviour. We included the biogenic amine depleting compound reserpine in view of previous studies that have implicated biogenic amine transmitters, in particular serotonin, in PPN motility and stylet thrusting [9]. The molecular mode of action lies in its potent inhibition of the vesicular monoamine transporter, VMAT [34] which elicits a global depletion of biogenic amines in the nervous system. Reserpine is one of the bioactive constituents in root extracts of *Rauwolfia serpentina*, commonly called Indian snakeroot, which have been used in herbal medicine for centuries for their sedative properties. Last century reserpine was one of the first drugs to be registered for the treatment of hypertension [35] and was also used as an antipsychotic [36]. We found that pre-treatment of *G. pallida* J2s with reserpine impaired their ability to invade the host root and this was accompanied by a reduction in motility which could contribute to this crop protecting effect. Cross-referencing the effective concentration of reserpine to the effective concentrations of the major nematicidal compounds e.g. aldicarb and abamectin, shows that reserpine is at least as, if not more, efficacious [37]. As reserpine is an inhibitor of the VMAT, which loads a range of biogenic amine neurotransmitters including serotonin and dopamine into their respective synaptic vesicular storage sites, the effect of reserpine on root invasion could be explained by depletion of any of these neurotransmitters.

To get a better insight into the neuropharmacological basis of the inhibitory action of reserpine on root invasion we developed a bioassay for monitoring stylet thrusting [10,38]. To assess the inhibitory effect of reserpine we first activated stylet thrusting by addition of either exogenous serotonin or fluoxetine. Serotonin induces rhythmic activity of the stylet and likely acts in a pathway that couples sensory perception of environmental cues to downstream signalling pathways to activate, or possibly disinhibit, stylet thrusting. This has parallels with *C. elegans* which also deploys serotonin signalling in response to environmental food cues to activate rhythmic pumping of its feeding organ, the pharynx [14,18]. We found that fluoxetine also induces stylet thrusting but in contrast to serotonin activation this depends on the availability of endogenous serotonin. Studies in *C. elegans* indicate this may be explained by an inhibition of serotonin reuptake [39] from the synapse and therefore increased levels of serotonin available to activate synaptic receptors. This approach therefore allowed us to test the hypothesis that reserpine exerts its effect by depleting endogenous serotonin. Interestingly, the concentration-response curve to fluoxetine was bell-shaped with lower stimulation observed at the highest concentrations tested. This may be explained by a direct receptor blocking effect of fluoxetine at high concentrations, a phenomenon also reported for *C. elegans* [11,39]. Notably, we found that reserpine exerted a selective and potent inhibition of stylet thrusting elicited by fluoxetine but not by serotonin indicating reserpine exerts its action by depleting endogenous serotonin. The estimated EC_50_for this effect of reserpine was 10 nM showing it has an extremely potent action, comparable to its effect on the mammalian VMAT1 and VMAT2 transporters [34]. Moreover, as might be predicted from a compound with this mode of action, the reserpine inhibition of fluoxetine-stimulated stylet thrusting was sustained and there was no recovery after 22 h removal from reserpine.

The serotonergic signalling pathway in PPNs is therefore a good candidate for new targets for crop protection. However, the molecular components of this have not previously been characterised despite the availability of PPN sequenced genomes [3,40]. In contrast, in *C. elegans* each element of the serotonergic pathway, from the synthetic enzymes through to the receptors has been identified and functionally assessed through the phenotypic delineation of their respective mutant strains. We made use of this to identify candidate orthologues of the respective *C. elegans* genes in the *G. pallida* genome. This revealed sequences encoding proteins with a greater than 35% amino acid identity to *C. elegans* VMAT [12], tryptophan hydroxylase [13], and the serotonin receptors SER-4 [24], SER-7 [15] and MOD-1 [27] (Table 1) which are therefore strong candidates as *G.pallida* functional orthologues of these key determinants of serotonergic signalling. In this study we focussed on further characterisation of SER-7 and MOD-1 given their known importance in *C. elegans* in coupling environmental food cues to behavioural outputs relating to foraging and feeding [15,27]. In *C. elegans* a number of these genes are subject to alternate splicing and this may also occur in *G. pallida* however for the purposes of this study we focused on cloning the longest variant of each transcript. Therefore we cannot discount the possibility that other, shorter, variants may exist.

We confirmed the functional role of each of the proteins encoded by the *G. pallida* genes identified from this bioinformatic screen for components of serotonergic pathways using a ‘model-hopping’ approach with *C. elegans*. For this we made use of *C. elegans* mutant strains carrying null mutations in each of the respective genes and for which there are characteristic behavioural or pharmacological phenotypes. We compared the ability of the wild-type *C. elegans* gene with that of the *G. pallida* putative orthologue to rescue the mutant phenotype. For each of the *G. pallida* genes we investigated, encoding VMAT, tryptophan hydroxylase, SER-7 and MOD-1, we were able to reinstate wild-type behaviour in the mutant. In effect, we recapitulated *G. pallida* serotonergic signalling in *C. elegans*.

We proceeded to investigate the relevance of each of these components of the serotonergic pathway to the impact of reserpine on root invasion behaviour. We have previously shown that RNAi has non-specific toxicity to *G. pallida* stylet thrusting which confounds the use of this experimental approach to assign gene function to root invasion behaviour [21]. Therefore we made use of the transgenic *C. elegans* strains expressing *G. pallida tph-1*, *ser-7* and *mod-1* to validate pharmacological blockers that could then be used to test for a role of each of these proteins in root invasion behaviour.

Wild-type *C. elegans* exposed to the selective tryptophan hydroxylase blocker CPA [22] showed a reduced rate of pharyngeal pumping in the presence of food, an effect which phenocopied the low pharyngeal pumping rate of the null *tph-1* mutant *mg280*. We observed a similar effect of CPA in *C. elegans* expressing *gpa-tph-1* thus indicating that CPA might be used as a pharmacological tool to interrogate the functional role of TPH in *G. pallida*. Indeed, exposure of *G. pallida* to CPA reduced fluoxetine-induced stylet thrusting and root invasion by J2s consistent with a pivotal role for serotonin signalling in these behaviours.

We found that methiothepin is a potent inhibitor of *G. pallida* stylet thrusting and root invasion. Given the very potent inhibition of stylet thrusting by methiothepin we also tested whether or not it would inhibit hatching of J2s from cysts as, like root invasion, this is also a stylet activity-dependent behaviour [5]. In contrast to the lack of effect of reserpine, methiothepin potently inhibited hatching. This discrepancy may be explained by a different ability of reserpine and methiothepin to diffuse across the protective cyst casing which can present a barrier to chemical access. The effect of methiothepin on hatching is intriguing as the J2s encased within the eggs still appear viable following the methiothepin exposure. We showed that both *G. pallida* SER-7 and MOD-1 are sensitive to methiothepin, similar to the *C. elegans* receptors [25,27], and these are strong candidates for mediating the inhibitory effect of methiothepin on PRD stimulated hatching, fluoxetine stimulated stylet thrusting and root invasion. However, methiothepin is known to act on other biogenic amine receptors, including SER-4 [41], and we cannot rule out a role for this receptor in particular. Further studies screening for selective antagonists of *G. pallida* SER-7 and MOD-1 has the potential to provide pharmacological tools that can more precisely address this question.

An outstanding question is where serotonin acts in the neural circuits of the worm to bring about a methiothepin-sensitive activation of stylet thrusting and root invasion. This behaviour is finely tuned to the worm’s environment and occurs at precise times during development triggered by cues from the rhizosphere or the plant which specifically engage sensory receptors and downstream signalling pathways to activate, or possibly disinhibit, stylet thrusting. By analogy with all other cue-dependent behaviours characterised in nematodes to date the circuit which controls PPN stylet thrusting will comprise morphologically specialised sensory neurons to detect environmental cues and will connect through interneurons that integrate signals from diverse inputs, which in turn output to motorneurons to directly regulate muscles controlling the stylet. There is a good precedent that comparison between *C. elegans* and other species of nematode provides insight into neuronal circuitry[42,43]. We found evidence for a pair of anterior serotonergic neurons in *G. pallida* that are in a location consistent with them performing a similar role to *C. elegans* anterior serotonergic neurons, either NSM, or ADF. Further studies are required to map these pathways in *G. pallida* and overlay the expression of serotonin receptors. In *C. elegans ser-7* and *mod-1* are widely expressed encompassing neurons, muscle and intestine consistent with the coordinating neurohormonal role of serotonin particularly with respect to feeding, foraging and energy homeostasis and it will be interesting to see whether a similar pattern plays out in *G. pallida* [26].

In conclusion, we have characterised molecular components of a serotonergic signalling pathway in *G. pallida* that is pivotal to root invasion behaviour and identify serotonin receptors that are candidate targets for crop protecting chemicals.

## Methods

### Globodera pallida maintenance and culture

Cysts of *G. pallida* (Pa2/3; population Lindley) were extracted from infested sand/loam following growth of host ‘Desiree’ potato plants. Dried cysts were treated with 0.1% malachite green solution for 30 mins followed by extensive washing in tap water. Cysts were then incubated in an antibiotic cocktail [44] at 4 °C overnight and washed five times with sterile tap water. To induce hatching, cysts were placed in a solution of 1 part potato root diffusate to 3 parts tap water. Root diffusate was obtained by soaking washed roots of three week-old potato plants in tap water at 4 °C overnight at a rate of 80g root/litre. The diffusate was filter-sterilised before use. Significant numbers of J2 typically began hatching 1 week after rehydration in the presence of potato root diffusate. Only J2 that had hatched within the previous 24 hours were used for experiments. Prior to the experiments the J2s were washed in ddH_2_O to remove potato root diffusate.

### Caenorhabditis elegans strains and culture

*C. elegans* were grown on Nematode Growth Medium (NGM) plates seeded with *Escherichia coli* (*OP50* strain) at 20ºC according to standard protocols [45]. N2 (Bristol strain) *C. elegans* were employed as wild-type. GR1321 is a strain carrying a deletion in *mg280* allele for *tph-1* (ZK1290.18) and a 9.8kb deletion in *vs166* for *cam-1*. *cam-1* encodes a receptor tyrosine kinase of the immunoglobulin superfamily. This strain displays 15 phenotypes and was used in the original studies on *C. elegans tph-1* gene characterisation [13]. Another strain, MT14984, contains a deletion only in the *tph-1* gene and displays one phenotype: reduced pharyngeal pumping (CGC, made by Dan Omura). However, this strain was not available to order from CGC at the time when the experiments were carried out. GR1321 was outcrossed with N2 *C. elegans* four times (CGC database) and additionally twice more in our laboratory. *ser-7(tm1325)* strain DA2100 carries a 742 bp deletion and 38 bp insertion and displays 5 phenotypes, one of which is reduced pharyngeal pumping in response to serotonin. It was outcrossed with N2 *C. elegans* ten times (CGC database). *cat-1* encodes a vesicular monoamine transporter (VMAT) and is an orthologue of human VMATs. *cat-1(ok411)X C. elegans* mutant strain RB681 has a 429 bp deletion and is predicted to be a functional null (Wormbase). MT9668 *mod-1* (*ok103*) *V* is a null allele due to the molecular nature of the mutation, covering the entire genomic sequence corresponding to the *mod-1* gene [27]. This strain was outcrossed with N2 *C. elegans* six times (CGC database). *mod-1* encodes a serotonin-gated chloride channel.

Transgenic *C. elegans* (described below) were always assayed in parallel with positive and negative controls for the pharyngeal pumping phenotype variants i.e. on the same day with N2 and mutant strain *C. elegans,* respectively.

### Drugs and Chemicals

Serotonin creatinine sulphate monohydrate (serotonin), methiothepin and 4-chloro-DL-phenylalanine methyl ester hydrochloride (CPA) were purchased from Sigma Aldrich (Dorset, UK). Fluoxetine hydrochloride was purchased from Enzo Life Sciences (Exeter, UK) and serpasil phosphate (reserpine) from Novartis (Surrey, UK). Agar plates containing compounds were prepared as follows: For serotonin creatinine sulphate monohydrate the compound was added to 60 ˚C NGM and stirred to give a 10 mM final concentration. The NGM was then used to pour plates. For fluoxetine and CPA the compounds were dissolved in M9. 500µl of each compound was evenly pipetted over the surface of a 6 cm plate containing 10 ml NGM to give the final desired concentration. The plates were allowed to dry for 2 hours before use. Serotonin and fluoxetine plates were stored at 4˚C for 1-2 weeks. Serpasil phosphate (reserpine) plates were prepared in a similar manner except the compound was dissolved in double distilled water and then diluted in HEPES buffer (pH 7.4). For stylet thrusting assays drugs were added to ddH_2_O (0.1% BSA) buffered to pH 7.4 to achieve the desired concentration. Methiothepin plates were prepared as described for the rest of the drugs except 10 µM dissolved in water was added to 10 ml of NGM. These drug plates were prepared freshly for use on the day of each experiment.

### Root invasion assays

J2s of *G. pallida* were first sterilised with 0.1% chlorhexidine digluconate, 0.5 mg/ml CTAB for 25 mins and washed three times with sterile 0.01% Tween-20. The J2s were then incubated with gentle agitation for 24 hours in water, 100 µM reserpine phosphate, 100 µM methiothepin or 100 µM CPA prior to inoculation of potato hairy root cultures with 25 J2s per infection point and 3 infection points per root system. Hairy root cultures were generated using *Agrobacterium rhizogenes* R1000 and multiplied as previously described [46]. Individual root systems of equivalent size growing on 9 cm plates containing Murashige and Skoog basal medium (Duchefa, Suffolk, UK) with 2% sucrose were selected for inoculation. Nine to 11 replicate plates were used for each treatment or control. Roots were stained with acid fuchsin 13 days after infection as described previously [47] and the number of nematodes in each root system counted. Each complete invasion assay was carried out on two separate occasions.

### G. pallida dispersal assays

Dispersal assays were performed on 5 cm plates filled with 10 ml of 2% agarose. For the reserpine assays worms were pre-soaked in either reserpine or water for 18 h. On the day of the assay 100 µl of potato root diffusate (PRD) was spread onto the plates and the plates were sealed with Parafilm. A grid with six equal concentric circles was placed under the assay plates and the centre was marked with a dot. Around 50-100 *G. pallida* J2s were pipetted with 5 µl of ddH 2O onto the centre (origin), the plate was re-sealed and the total number of worms was counted as soon as the liquid had absorbed. The number of J2s in each concentric circle was counted after 1 and 2 h.

### G. pallida stylet thrusting assays

Stylet thrusting assays were conducted in 20 mM HEPES pH 7.4. J2s were pipetted into 3 ml of the test solutions in 30 mm Petri dishes, mounted on the stage of an inverted microscope for viewing and the number of stylet thrusts per minute was counted at various time points as stated in figure legends. A single movement of the stylet knob forwards and then backwards to its original position was counted as one stylet thrust. Control assays were conducted in the presence of either 20 mM HEPES alone or 20 mM HEPES with drug vehicle. All assays were conducted at room temperature (20-22°C). All drug solutions were made up on the day of use. Each solution contained 0.01% bovine serum albumin to prevent J2s from sticking to the Petri dish. For experiments in which J2s were pre-treated with reserpine phosphate or CPA, the worms were pre-soaked in the drug solutions for 24 hours.

### Structural characterisation of C. elegans and G. pallida tph-1, ser-7, cat-1 and mod-1

*G. pallida* putative orthologues of *C. elegans tph-1* (WBGene00006600), *ser-7* (WBGene00004780), *cat-1* (WBGene00000295) and *mod-1* (WBGene00003386) were first identified by BLASTP searches of the predicted protein dataset (May 2012) at http://www.sanger.ac.uk/cgi-bin/blast/submitblast/g_pallida followed by reciprocal searches of the *C. elegans* protein set at wormbase.org. [4]. Each predicted *G. pallida* gene was located in the draft genome assembly after a BLASTN search of the scaffolds via the *G. pallida* BLAST server as above. The predicted gene models were manually assessed for concordance with the mapped transcripts and primers were designed to amplify the complete coding regions.

The primary amino acid sequences of *C. elegans* and *G. pallida* CAT-1, TPH-1, SER-7 and MOD-1 proteins were aligned using the UNIPROT Clustal Omega program. Clustal-Omega uses the HHalign algorithm and its default settings as its core alignment engine [49].

### Cloning of C. elegans and G. pallida tph-1, ser-7, cat-1 and mod-1

RNA was extracted from mixed stages *C. elegans* and from J2 *G. pallida* using an RNeasy Mini Kit (Qiagen, UK) according to the manufacturer’s instructions. cDNA was reverse transcribed from 200 ng (*C. elegans*) or 500 ng (*G. pallida*) of total RNA using Superscript III reverse transcriptase with oligo-dT primers (Life Technologies, UK). *C. elegans tph-1a*, *ser-7a* and *mod-1a* were then amplified by PCR from 2 µl of cDNA using proof reading Phusion DNA polymerase (Thermo Scientific, UK), followed by addition of 1 µl of Taq DNA polymerase (Promega, UK), and continued with 10 min at 72 °C. *G. pallida* predicted orthologues were amplified from 1 µl of cDNA using Platinum HiFi polymerase (Life Technologies, UK). PCR products were analysed by agarose gel electrophoresis and cloned into pCR8/GW/TOPO vector according to the manufacturer’s instructions (Life Technologies, UK). TOP10 chemically competent cells (Life Technologies, UK) were transformed with TOPO reaction and plated onto spectinomycin (100 mg/ml) selective plates overnight at 37°C. *C. elegans cat-1* cDNA in the pDONR201 vector (W01C8.6) was purchased from the *C. elegans* library (GE Healthcare Dharmacon, Thermo Fisher Scientific Biosciences, GmbH) and used as template to amplify and clone *cat-1a* as described for the other genes. The orientation of the cloned genes was confirmed by digests with restriction enzymes. Subsequently, colonies that contained genes in 5’ to 3’ orientation were sequenced by a commercial company (Eurofins Genomics, UK). All *C. elegans* sequences subsequently used were in agreement with published data. Out of 7 sequenced colonies, 1 colony was selected as it contained a complete sequence for *cat-1a* with correct “start” and “stop” codons.

The *C. elegans Psnb-1* promoter was digested from the pBK3.1 vector [48] with HindIII and XbaI enzymes. pDEST vector (Life Technologies, UK) was digested with the same restriction enzymes and the fragments for *Psnb-1* promoter and linear pDEST vector were purified out of the agarose gel. The *Psnb-1* promoter was ligated into the pDEST vector in a 3:1 ratio with T4 DNA ligase (Promega, UK) overnight at 4 °C and transformed into One Shot ccdB Survival 2 T1R Competent Cells (Life Technologies) grown on ampicillin (100 µg/µl) and chloramphenicol (30 µg/ml) selective plates. DNA insertion was confirmed by restriction digests and clones sequenced by Eurofins Genomics, UK. All sequenced vectors contained the predicted sequence for *Psnb-1*.

The *Pmod-1 C*. *elegans* promoter was amplified from genomic DNA (Life Technologies, UK), was digested with both NheI and KpnI restriction enzymes, and the similarly digested *mod-1* promoter fragment was ligated into pDEST vector following the same protocol described above.

The *in vitro* recombination between an entry pCR8/GW/TOPO clone containing *C. elegans* or *G. pallida tph-1*, *ser-7, cat-1, mod-1* and a destination vector pDEST;*Psnb-1* or pDEST;*Pmod-1* as indicated, was performed using Gateway^®^ LR Clonase^®^ II enzyme mix (Life Technologies, UK) according to the manufacturer’s instructions to generate expression clones for *C. elegans* microinjections. The resulting plasmids were propagated in TOP10 cells grown on ampicillin selective plates (100 µg/ml) and confirmed by restriction digests and sequencing.

### C. elegans transgenic experiments

*tph-1(mg280)*, *ser-7(tm1325)* and *cat-1(ok411) C. elegans* were injected with plasmids to drive expression of either *C. elegans* or *G. pallida tph-1*, *ser-7* or *cat-1* from a synaptobrevin promoter, *Psnb-1*, which drives expression in all neurons. *mod-1(ok103) C. elegans* were injected with a plasmid to drive expression of either *C. elegans* or *G. pallida mod-1* from the native promoter *Pmod-1*. The plasmids were injected at 30 ng µl^−1^, except for *Pmod-1* which was injected at 10 ng µl^−1^. Transformed worms were identified by co-injecting L3785 (*Pmyo-3::gfp*) plasmid (50 ng µl^−1^) (a gift from Andrew Fire), which drives expression of green fluorescent protein (GFP) from the body wall muscle promoter *Pmyo-3* [50]. The co-injected *gfp* transformation marker forms an extra-chromosomal array with the plasmids carrying the gene sequence and thus worms with fluorescent green body wall muscle can be identified as carrying the plasmid of interest. For all the experiments, at least two independently transformed stable lines of transgenic *C. elegans* expressing *C. elegans* or *G. pallida tph-1*, *ser-7, cat-1* or *mod-1* were assayed. Results for the independent lines for each construct were in good agreement and the data presented are the pooled data from these independent lines.

### Pharmacological characterisation of the transgenic C. elegans strains in pharyngeal pumping assays

Experiments were performed on one day old age synchronised worms by picking L4 stage a day before the assay. Due to the translucent nature of the worm, pharyngeal pumping may be scored in the intact animals by counting the movements of the grinder in the terminal bulb: one complete up and down motion is counted as a single pharyngeal pump. The number of pharyngeal pumps was counted on *E. coli* OP50 in N2, in the mutants *tph-1 (mg280)*, *ser-7 (tm1325)*, *cat-1 (ok411)*, *mod-1(ok103)* and in mutants with ectopic expression of *C. elegans* (*ce*) or *G. pallida* (*gpa*) *tph-1*, *ser-7* or *cat-1* in all of the neurons (*Psnb-1* promoter).

To test the effects of CPA (4-chloro-DL-phenylalanine methyl ester hydrochloride) on pharyngeal pumping one day old adult *C. elegans* for N2, *tph-1 (mg280)* and *tph-1-1 (mg280)* with ectopic expression of *C. elegans* (*ce*) or *G. pallida* (*gpa*) *tph-1* were placed onto *E. coli* OP50 on NGM plates containing CPA and the number of pharyngeal pumps on food per min was scored after 2 and 18 hours. Pharyngeal pumping of *ser-7* mutants expressing either *C. elegans* or *G. pallida ser-7* was tested after 20 min exposure to 10 mM serotonin in the absence of food. Additionally, an intact pharynx containing a terminal bulb was dissected from the rest of the worm with a razor blade. The pharyngeal preparation was placed into 3 ml of Dent’s saline (140 mM NaCl, 10 mM HEPES, 10 mM D-glucose, 6 mM KCl, 3 mM CaCl_2_, 1 mM MgCl_2_, pH 7.4 with 1 mM NaOH with 0.1% BSA) and the drugs serotonin or methiothepin) at a range of concentrations as indicated, in a 25 mm Petri dish. Pharyngeal pumps were scored visually, as for the intact worms, for 1 min. For controls, the pharyngeal pumping was scored in Dent’s saline (0.1% BSA) without drugs with appropriate vehicle controls. To test the effects of fluoxetine one day old *C. elegans* for N2, *cat-1 (ok411)* and *cat-1 (ok411)* with ectopic expression of *C. elegans* (*ce*) or *G. pallida* (*gpa*) *cat-1* were placed onto unseeded NGM plates containing fluoxetine at a range of concentrations as indicated and the number of pharyngeal pumps per min was scored after 1 h.

### Pharmacological characterisation of the transgenic C. elegans strains in thrashing paralysis assay

In order to study the functionality of both *C. elegans* and *G. pallida* MOD-1 channels we examined the pharmacological response of MOD-1 to serotonin, using a thrashing paralysis assay as described by Ranganathan et al [27]. Briefly, 10 to 20 animals (L4+1 day stage) were placed in 200 µl of 33 mM serotonin dissolved in M9 buffer in 96-well microtitre wells. The serotonin resistance was scored, observing the swimming behaviour, every minute for a total time of 20 min. An animal was considered immobile if it did not exhibit any swimming motion for a period of 5 s.

To test whether methiothepin was able to block the serotonin-induced paralysis, we carried out the thrashing paralysis assay as described above, but with a pre-incubation with or without 10 µM methiothepin for 120 min on an NGM plate with *E. coli* OP50 bacteria before placing the worms in the wells of the microtitre plate with serotonin. For the methiothepin treatment group, 10 µM methiothepin was included in the wells of the microtitre plate. Wild type (N2), *mod-1* (*ok103*), and the transgenic strains expressing *gpa-mod-1* in *mod-1(ok103)* were tested.

### G. pallida hatching assays

*G. pallida* cysts were washed in ddH_2_O and individual cysts were transferred to wells in a 24 well plate, containing tap water 1:3 PRD or drug solutions made using 1:3 PRD. A vehicle control was performed for each experiment. Methiothepin was dissolved in 100% ethanol and were added to a solution of 1:3 PRD to give a final ethanol concentration of 0.5%. The cysts were soaked in 1:3 PRD solution in the presence of drug, methiothepin or reserpine, for up to 25 days. During this period hatched J2s were counted and removed from the wells. The solution in which the cysts were soaked was replaced each time a count was taken. The cysts were then removed from the drug solution and transferred to the wells of another 24 well plate containing 1:3 PRD alone to assess hatching recovery. J2 hatching was then counted in the same manner. Throughout each individual experiment the same PRD batch was used. Cumulative hatch of J2s per cyst was plotted over time. At the conclusion of the hatching experiment, cysts were transferred to ddH_2_O and cracked open with a razor to count the number of unhatched eggs per cyst.

### *In situ* hybridization

Digoxygenin (DIG)-labelled strand-specific DNA probes for *gpa-tph-1* were prepared from a 142 bp template cDNA fragment amplified with the primers 5’-AACGGCCGATGTACTGCTAG-3’ (F) and 5’-CGACTCTGTCCGGGTCAAAA-3’ (R) according to a standard protocol [32].J2 *G. pallida* were fixed, permeabilised and hybridised with the probes as described and NBT/BCIP was used for probe detection following incubation with an anti-DIG, alkaline phosphatase-conjugated antibody.

### Statistical analysis

Data points in graphs are presented as the mean ± standard error of the mean for the number of experiments as shown in individual figures. Data were plotted using GraphPad Prism 7.01software (San Diego, California). Statistical significance was determined either by unpaired Student’s t-test, one-way or two-way ANOVA as appropriate; significance level set at P< 0.05, followed by Bonferroni multiple comparisons as appropriate. EC_50_ values with 95% confidence intervals were determined using GraphPad Prism 7.01 by plotting log concentration agonist against response and fitting the data to the equation; Y=Bottom + (Top-Bottom)/(1+10^((LogEC50-X))).

#### Acknowledgements

Anna Crisford and Elizabeth Ludlow were supported by Biotechnology and Biological Sciences (BBSRC) grant number BB/J006890/1. Fernando Calahorro was supported by an award from Bayer Grants4Targets. Jennifer Hibbard was supported by BBSRC grant number BB/J006017/1. Some *C. elegans* strains were provided by the CGC, which is funded by NIH Office of Research Infrastructure Programs (P40 OD010440). James Dillon (University of Southampton, UK) provided the image of *C. elegans* NSM neurons.

**Conflict of Interest Statement:** Holden-Dye, O’Connor and Urwin are inventors for filed UK Patent Application no. 1710057.9 Reserpine Seed Coating.

## References

1. Chitwood D (2003) Research on plant-parasitic nematode biology conducted by the United States Department of Agriculture-Agricultural Research Services. Pest Manag Sci 59.

2. Holden-Dye L, Walker RJ (2014) Anthelmintic drugs and nematicides: studies in Caenorhabditis elegans The *C elegans* Research Community, WormBook, doi/10.1895/wormbook.1.143.2.

3. Eves-van den Akker S, Laetsch DR, Thorpe P, Lilley CJ, Danchin EGJ, et al. (2016) The genome of the yellow potato cyst nematode, *Globodera rostochiensis*, reveals insights into the basis of parasitism and virulence. Genome Biol 17: 124.

4. Cotton JA, Lilley CJ, Jones LM, Kikuchi T, Reid AJ, et al. (2014) The genome and life-stage specific transcriptomes of *Globodera pallida* elucidate key aspects of plant parasitism by a cyst nematode. Genome Biol 15: R43–R43.

5. Perry RN, Moens M (2013) Plant Nematology: Cabi.

6. Trudgill DL (1991) Resistance to and tolerance of plant parasitic nematodes in plants. Ann Rev Phytopathol 29: 167–192.

7. Rolfe RN, Perry RN (2001) Electropharyngeograms and stylet activity of second stage juveniles of *Globodera rostochiensis*. Nematology 3: 31–34.

8. Holden-Dye L, Walker R (2011) Neurobiology of plant parasitic nematodes. Inv Neurosci: 1–11.

9. Masler E (2007) Responses of *Heterodera glycines* and *Meloidogyne incognita* to exogenously applied neuromodulators. J Helminthol 81: 421–427.

10. Kearn J (2015) Mode of action studies on the nematicide fluensulfone. PhD Thesis University of Southampton.

11. Kullyev A, Dempsey CM, Miller S, Kuan C-J, Hapiak VM, et al. (2010) A genetic survey of fluoxetine action on synaptic transmission in *Caenorhabditis elegans*. Genetics 186: 929–941.

12. Duerr JS, Frisby DL, Gaskin J, Duke A, Asermely K, et al. (1999) The *cat-1* gene of Caenorhabditis elegans encodes a vesicular monoamine transporter required for specific monoamine-dependent behaviors. J Neurosci 19: 72–84.

13. Sze JY, Victor M, Loer C, Shi Y, Ruvkun G (2000) Food and metabolic signalling defects in a *Caenorhabditis elegans* serotonin-synthesis mutant. Nature 403: 560–564.

14. Niacaris T, Avery L (2003) Serotonin regulates repolarization of the *C. elegans* pharyngeal muscle. J Exp Biol 206: 223–231.

15. Hobson R, Hapiak V, Xiao H, Buehrer K, Komuniecki P, et al. (2006) SER-7, a *Caenorhabditis elegans* 5-HT7-like receptor, is essential for the 5-HT stimulation of pharyngeal pumping and egg laying. Genetics 172: 159–169.

16. Rogers C, Franks C, Walker R, Burke J, Holden-Dye L (2001) Regulation of the pharynx of *Caenorhabditis elegans* by 5-HT, octopamine, and FMRFamide-like neuropeptides. J Neurobiol 49: 235 – 244.

17. Nonet M, Saifee O, Zhao H, Rand J, Wei L (1998) Synaptic transmission deficits in Caenorhabditis elegans synaptobrevin mutants. Journal of Neuroscience 18: 70 – 80.

18. Song B, Avery L (2012) Serotonin activates overall feeding by activating two separate neural pathways in *Caenorhabditis elegans*. J Neurosci 32: 1920–1931.

19. Ranganathan R, Sawin ER, Trent C, Horvitz HR (2001) Mutations in the Caenorhabditis elegans serotonin reuptake transporter MOD-5 reveal serotonin-dependent and -independent activities of fluoxetine. J Neurosci 21: 5871–5884.

20. Dallière N, Bhatla N, Luedtke Z, Ma DK, Woolman J, et al. (2015) Multiple excitatory and inhibitory neural signals converge to fine-tune *Caenorhabditis elegans* feeding to food availability. FASEB J 30: 836–848.

21. Crisford A, Ludlow E, Lilley C, Kearn J, Urwin P, et al. (2016) The effect of double-stranded RNA on stylet behaviour in *Globodera pallida*. Aspects Appl Biol 130, 4th Symposium of Potato Cyst Nematode Management (including other nematode parasites of potatoes),: pp. 1–5.

22. Jequier E, Lovenberg W, Sjoerdsma A (1967) Tryptophan hydroxylase inhibition: the mechanism by which p-chlorophenylalanine depletes rat brain serotonin. Mol Pharmacol 3: 274–278.

23. Carnell L, Illi J, Hong S, McIntire S (2005) The G-protein-coupled serotonin receptor SER-1 regulates egg laying and male mating behaviors in Caenorhabditis elegans. J Neurosci 25: 10671 – 10681.

24. Harris GP, Hapiak VM, Wragg RT, Miller SB, Hughes LJ, et al. (2009) Three distinct amine receptors operating at different levels within the locomotory circuit are each essential for the serotonergic modulation of chemosensation in *Caenorhabditis elegans*. J Neurosci 29: 1446–1456.

25. Hobson RJ, Geng J, Gray AD, Komuniecki RW (2003) SER-7b, a constitutively active Gαs coupled 5-HT7-like receptor expressed in the *Caenorhabditis elegans* M4 pharyngeal motorneuron. J Neurochem 87: 22–29.

26. Komuniecki RW, Hobson RJ, Rex EB, Hapiak VM, Komuniecki PR (2004) Biogenic amine receptors in parasitic nematodes: what can be learned from *Caenorhabditis elegans*? Mol Biochem Parasitol 137: 1–11.

27. Ranganathan R, Cannon S, Horvitz H (2000) MOD-1 is a serotonin-gated chloride channel that modulates locomotory behaviour in C. elegans. Nature 408: 470 – 475.

28. Jones AK, Sattelle DB (2008) The cys-loop ligand-gated ion channel gene superfamily of the nematode, *Caenorhabditis elegans*. Inv Neurosci 8: 41–47.

29. Flavell Steven W, Pokala N, Macosko Evan Z, Albrecht Dirk R, Larsch J, et al. (2013) Serotonin and the neuropeptide PDF initiate and extend opposing behavioral states in *C. elegans*. Cell 154: 1023–1035.

30. Law W, Wuescher LM, Ortega A, Hapiak VM, Komuniecki PR, et al. (2015) Heterologous expression in remodeled *C. elegans*: A platform for monoaminergic agonist identification and anthelmintic screening. PLOS Pathog 11: e1004794.

31. Pemberton D, Franks C, Walker R, Holden-Dye L (2001) Characterization of glutamate-gated chloride channels in the pharynx of wild-type and mutant *Caenorhabditis elegans* delineates the role of the subunit GluCl-alpha2 in the function of the native receptor. Mol Pharmacol 59: 1037–1043.

32. Thorpe P, Mantelin S, Cock PJA, Blok VC, Coke MC, et al. (2014) Genomic characterisation of the effector complement of the potato cyst nematode Globodera pallida. BMC Genomics 15: 923.

33. Guo X, Wang J, Gardner M, Fukuda H, Kondo Y, et al. (2017) Identification of cyst nematode B-type CLE peptides and modulation of the vascular stem cell pathway for feeding cell formation. PLOS Pathog 13: e1006142.

34. Erickson JD, Eiden LE, Hoffman BJ (1992) Expression cloning of a reserpine-sensitive vesicular monoamine transporter. Proc Natl Acad Sci USA 89: 10993–10997.

35. Trapold JH, Plummer AJ, Yonkman FF (1954) Cardiovascular and respiratory effects of Serpasil, a new crystalline alkaloid from the *Rauwolfia serpentina* benth, in the dog. J Pharm ExpTherap 110: 205–214.

36. Noce RH, Williams DB, Rapaport W (1954) Reserpine (serpasil) in the management of the mentally ill. J Am Med Assoc 158: 11–15.

37. Faske TR, Starr JL (2006) Sensitivity of *Meloidogyne incognita* and *Rotylenchulus reniformis* to Abamectin. J Nematol 38: 240–244.

38. Kearn J, Lilley C, Urwin P, O Connor V, Holden-Dye L (2017) Progressive metabolic impairment underlies the novel nematicidal action of fluensulfone on the potato cyst nematode *Globodera pallida*. Pest Biochem Physiol in press.

39. Dempsey CM, Mackenzie SM, Gargus A, Blanco G, Sze JY (2005) Serotonin (5HT), fluoxetine, imipramine and dopamine target distinct 5HT receptor signaling to modulate *Caenorhabditis elegans* egg-laying behavior. Genetics 169: 1425–1436.

40. Opperman CH, Bird DM, Williamson VM, Rokhsar DS, Burke M, et al. (2008) Sequence and genetic map of Meloidogyne hapla: A compact nematode genome for plant parasitism. Proceedings of the National Academy of Sciences 105: 14802–14807.

41. Petrascheck M, Ye X, Buck LB (2007) An antidepressant that extends lifespan in adult Caenorhabditis elegans. Nature 450: 553–556.

42. Walrond JP, Kass IS, Stretton AO, Donmoyer JE (1985) Identification of excitatory and inhibitory motoneurons in the nematode Ascaris by electrophysiological techniques. J Neurosci 5: 1–8.

43. Bumbarger Daniel J, Riebesell M, Rödelsperger C, Sommer Ralf J (2012) System-wide rewiring underlies behavioral differences in predatory and bacterial-feeding nematodes. Cell 152: 109–119.

44. Urwin PE, Lilley CJ, McPherson MJ, Atkinson HJ (1997) Resistance to both cyst and root-knot nematodes conferred by transgenic *Arabidopsis* expressing a modified plant cystatin. Plant J 12: 455–461.

45. Brenner S (1974) The genetics of *Caenorhabditis elegans*. Genetics 77: 71–94.

46. Urwin PE, Atkinson HJ, Waller DA, McPherson MJ (1995) Engineered oryzacystatin-I expressed in transgenic hairy roots confers resistance to *Globodera pallida*. Plant J 8: 121–131.

47. Atkinson HJ, Urwin PE, Clarke MC, McPherson MJ (1996) Image analysis of the growth of Globodera pallida and Meloidogyne incognita on transgenic tomato roots expressing cystatins. J Nematol 28: 209–215.

48. Wang Z-W, Saifee O, Nonet ML, Salkoff L (2001) SLO-1 potassium channels control quantal content of neurotransmitter release at the *C. elegans* neuromuscular junction. Neuron 32: 867–881.

49. Soding J (2005) Protein homology detection by HMM-HMM comparison. Bioinformatics 21: 951–960.

50. Okkema PG, Harrison SW, Plunger V, Aryana A, Fire A (1993) Sequence requirements for myosin gene expression and regulation in *Caenorhabditis elegans*. Genetics 135: 385–404.

51. Eves-van den Akker S, Lilley CJ, Jones JT, Urwin PE (2014) Identification and Characterisation of a Hyper-Variable Apoplastic Effector Gene Family of the Potato Cyst Nematodes. PLoS Pathog 10: e1004391.

52. Dillon J, Franks CJ, Murray C, Edwards RJ, Calahorro F, et al. (2015) Metabotropic glutamate receptors: modulators of context-dependent feeding behaviour in *C. elegans*. J Biol Chem.

53. Jafari G, Xie Y, Kullyev A, Liang B, Sze JY (2011) Regulation of extrasynaptic 5-HT by serotonin reuptake transporter function in 5-HT-absorbing neurons underscores adaptation behavior in *Caenorhabditis elegans*. J Neurosci 31: 8948–8957.

